# Population Structure and Ecology of the African elephant (*Loxodonta africana*, Blumenbach, 1797) in Chebera Churchura National Park, Ethiopia

**DOI:** 10.1101/2022.07.14.500075

**Authors:** Adane Tsegaye, Afework Bekele, Anagaw Atikem

**Author notes:** Corresponding author: Adane Tsegaye, Email =, Telephone: **.

## Abstract

An investigation on population structure and ecology of the African elephants (*Loxodonta africana*) was carried out in Chebera Churchura National Park, Ethiopia during the wet and dry seasons of 2020 –2021. Sample counts using distance sampling of the African elephants were carried out in an area of 1, 410 km^2^. The total population estimated was 756 individuals, and the mean population density estimated was 0.53/km^2^. Among these, females constituted 52.12% and males 36.07%. The remaining 11.8% of the population was young of both sexes. It was difficult to categorize the young into male and female in the field, as their primary sexual characteristics were not easily visible.

Male to female sex ratio was 1.00:-1.42. Age structure was dominated by adults, which constituted 53.83% of the total population. Sub-adults comprised 19.11%, juveniles contributed 15.18% and calves accounted for 12.11% of the population. The herd size ranged from 1 to 149 individuals and the mean herd size during wet and dry seasons were 16.5 and 50.25, respectively. The African elephants were distributed in four habitat types: grassland, woodland, montane forest and riverine habitats in the study area. They were observed more in the riverine vegetation types during the dry season. Relative abundance of food resources, green vegetation cover and water availability in the area were the major factors governing their distribution in the present study area.

## 1. Introduction

African elephants were widely distributed across the continent prior to colonial times and were widespread in all over sub-Saharan Africa in diverse habitats ranging from tropical forests to semi-arid bushlands and desert. However, at present they have highly reduced both in number and range, due to human induced factors mainly poaching for ivory and habitat loss and fragmentation. At present African elephants occur only in 38 range States. In Africa, the African elephant is the only surviving species from order Probocidea. Currently, two subspecies of African elephants are recognized: the savanna elephant and the forest elephant. Currently, they are treated as separate species by IUCN and listed the savanna elephant, *Loxodonta africana africana* as “endangered” and the forest elephant, *Loxodonta africana cyclotis* as “crtically endangered” species due to the huge decline both in their number and range (IUCN, 2021).

Ethiopia is one of the African countries that harbours elephants (Blanc *et al*., 2003; Admasu 2006; Demeke, 2010). However, well-known historical information about elephants for Ethiopia before the 1960s is lacking. In the 1970s the status of elephants in Ethiopia was estimated to number from 6000 to 10,000 (Demeke, 2009; Demeke, 2010). However, intensive poaching, habitat loss and fragmentation resulted in approximate loss of 90% of the total elephant with total extirpation from 6 of 16 sites since 1980s (Demeke, 2010). The total national population is currently estimated at between ∼1850-1900 animals occurring in 6 main populations of Omo, Mago, Gambella, Kafta-Sheraro, Chebera Churchura National Parks and the Babille Elephant Sanctuary (EWCA, 2015).

Detailed information on population structure and status are critical for establishing and predicting population trends of keystone species as well as informing management decisions. As a key stone species, elephants are noted for the capacity to modify their habitats, intelligence, close family ties, social complexity and key players in structuring entire habitats. Their extinction at local or large scale level definitely will have cascading effects (Jyoti *et al*., 2020).

Appropriate management decisions and practices of any conservation measure towards a species or habitat requires a scientific investigation (ecological study) of the target species with a reliable estimate of population size, structure and the habitat where it lives (Kumara1 *et al*., 2012; Mekonen, 2019). In order to evaluate the conservation status or understand the vulnerability to different threats on a species it is vital to know the population size, distribution, its geographic range and the rate of population decline as well as habitat preferences (Motsumi *et al*., 2007). But the above information was lacking on the threatened African Elephant population in Chebra Churchura National Park Ethiopia, where it is crucial to design a biologically sound conservation strategy.

Chebera Churchura National Park (CCNP) is home to almost one third of the national elephant population with a high diversity of flora and fauna. However, except for survey on the diversity of small mammals and few outdated surveys of larger mammal and bird diversity, there is inadequate recent information about the population structure and status of African elephant in the area. As human threats continue to impact on natural habitats, there is an increasing need to regularly monitor the status and trends in large vertebrate populations and their habitats which need a baseline data. Thus assessment of the current population and distribution of the African elephant in and around CCNP will give us full information about the animal in the study area. The National Park thus provides baseline information vital for the selection, establishment, designing and implementation of the management of the species.

Accurate estimation of elephant population needs appropriate scientific methods. A line transect method involving distance sampling (Burnham *et al*., 1980) will be more appropriate technique for assessing elephant population in protected areas that have closed canopy vegetation cover like CCNP where aerial survey is difficult to implement (Jyoti *et al*., 2020).

## 2. The Study Area and Methods

### 2.1. Study Area

Chebera Churchura National Park (CCNP) is located in the southwestern part of Ethiopia, in the newly established South Western Ethiopian Administrative Region. The Park is located between Dawro and Konta Zones. It covers an area of 1410 km^2^ and lies between the coordinates 36° 27’00’’-36° 57’14’’E and 6°56’05’’-7° 08’02’’N (Tsegaye *et al*., 2015) (Fig 1).

**Figure 1.**
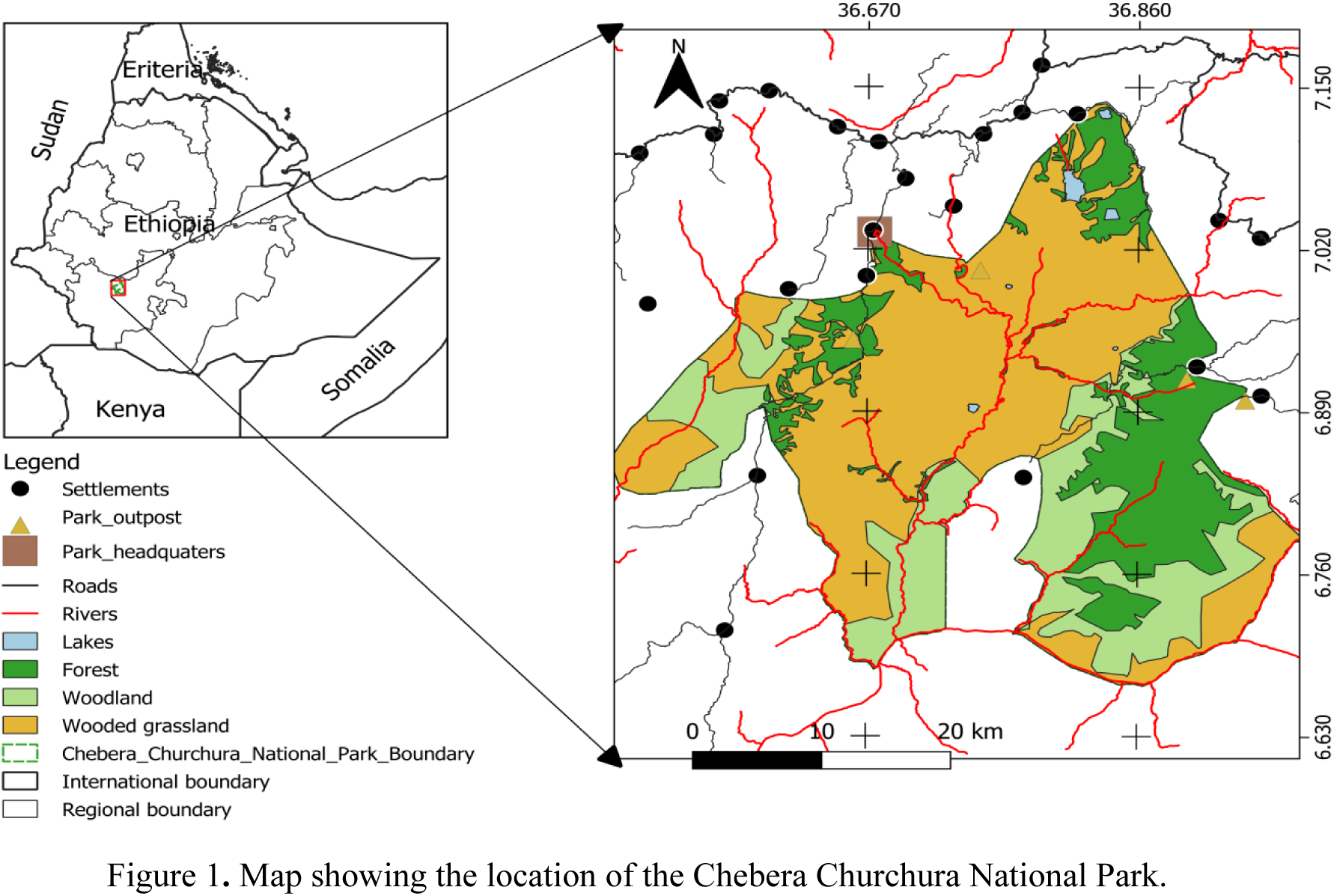
Map showing the location of the Chebera Churchura National Park.

Chebera Churchura is characterized by relatively hot climatic conditions. The rainfall distribution is unimodal. The average amount of annual rainfall in the area varies from 1000 to 3500 mm. The area has a uniform and long rainfall season (between March and September with a peak in July). The dry season is from November to February, with mean maximum temperatures varying between 27 and 29oC. The hottest months are January and February while, the coldest months are July and August with the mean maximum and minimum temperatures of 28oC and 12oC, respectively (Weldeyohanes, 2006).

The vegetation cover of the area is categorized as follows: wooded grassland, woodland, montane forest and riparian forest. Wooded grassland accounts for 55.6% of the study area. It covers most of the undulating landscapes above the floor of the valleys and gorges. Although the grass species show local variation, the dominant grass species is the elephant grass *Pennisetum* sp (Tsegaye *et al*., 2015).

The tree species are deciduous and include *Combretum* sp. in association with *Terminalia albiza*. Woodland habitat covers about 13.2% of the total area while the riparian forest habitat covers only 3% of the total area of the Park. The montane forest habitat covers about 27.2% of the total area of the park. Dominant tree species include *Juniperus procera, Podocarpus falcatus* and other broad leaved species (Woldeyohannes, 2006). Montane forest vegetation occurs in the eastern and northwestern highlands of the study area. It is dominated by tree species and characterized by a crown cover of up to 50%, with multistoried structure. Climbers and saprophytes are important floristic components of the habitat. Riparian forest covers about (4%) along the course of the rivers. These are Zigna, Shoshima, Wala, Tikurwuha, Mensa, Oma and other small seasonal tributaries. This habitat is characterized by mixed vegetation type composed of large trees and herbaceous species. Dominant plant species in this habitat are *Ficus* sp., *Phoenix* sp., Costa sp., *Albizia grandibracteata, Chionanthus mildbraedii, Grewia ferruginea, Aspilia mossambicensis, Arundo donax* and *Ehretia cymosa* (Timer, 2005; Admasu, 2006).

The principal ethnic groups found around CCNP are Dawro and Konta Nationalities. Other minority groups include Tsara, Menja, Mena and Bacha. Dawro ethnic group which inhabit the eastern highland and few areas of the southeastern lowland areas. These people do not make extensive use of the lowlands except along the periphery. Konta ethnic group occupies the north and northwestern highland areas (Datiko, 2013).

Mixed agricultural practices are the main livelihood of the majority of the inhabitants living around the Park. The people practice traditional agricultural systems that combine perennial and annual cultivation with livestock rearing (Datiko, 2013). The minority groups of people also lead their livelihood by collecting and selling wild honey, spices, wild coffee and edible roots from the forests of some of the wild plants (Tsegaye *et al*., 2015).

### 2.2. Methods

Based on the topography and habitat types the total area of the study was classified into smaller sample units and each of the sample area represent one or more of the major habitats of the study area. The number of transects that was laid on the major habitats were based on the total size of the area, topography and vegetation type. The representative transects that cross each habitat was randomly selected representing all of the major habitat types.

By modifying the unequally sized sample unit ratio following Norton Griffiths, (1978) ; Ndhlovu and Balakrishnan, (1991); Megaze (2015) and (Jyoti *et al*., 2020) were adopted for sampling design. Out of the total blocks of the study area a number of representative sample blocks were randomly selected. The sampling blocks selected from each habitat type represent 25% of each of the surveyed areas. Randomly selected transects were then established in each block. A total of 41 transects were sampled. The number of transects in each of the census zones varied depending on visibility, topography and size (Norton-Griffiths, 1978; Ndhlovu and Balakrishnan, 1991; Jyoti *et al*., 2020). Survey was conducted using subsidiary tracks guided by GPS and compass in each randomly selected block along selected transects. The length of transects varied from 4.5 to 5 km. Each of these transect-lines was located randomly in the study area in each of the habitat type using global positioning system. Adjacent transects were 2000 to 2500 m apart. All transects were roughly parallel to each other and their ends we’re not less than 1000 m far from the habitat edge. Special care was taken to keep each consecutive transect at a distance of around 2–2.5 km away from each other (Koster and Hart, 1998).

Transects for observation were laid based on the four major vegetation categories of the study area as follow:

1. Census zone 1 (Wooded grassland): This habitat type covered an estimated area of 784 km^2^ in CCNP. Total area sampled was 196 km^2^. A total of 12 transects of different length were laid in this habitat.
2. Census zone 2 (Woodland): This habitat type covered an estimated extent of 186 km^2^ area in CCNP. Total area sampled was 46.5 km^2^. There were 10 transects of different lengths laid in this habitat.
3. Census zone 3 (Montane forest): This habitat type covered an area of 384 km^2^. Total area sampled was 96 km^2^ where 11 transect of different length were laid.
4. Census zone 4 (Riparian forest): This habitat type covered an area of 56 km^2^. Total area sampled was 196 km^2^. Total area sampled was 14 km^2^. There were 8 transects of different length laid in this habitat.

### 2.3. Field Study

The first step prior to the actual field work was a reconaissance survey. It was conducted by vehicle and on foot during May 2019. Reconaissance observations were carried out to provide information on accessibility, climate, vegetation cover, topography, infrastructure, fauna, distribution of African elephant and launching sampling plans. Data collection was carried out during both wet and dry seasons between the years 2020 to 2021. Seasonal differences in the population size, distribution and habitat association of African elephants were recorded and compared.

Separation of wet and dryseasons was based on the change of rainfall pattern. Quantitative data were collected on the population size, age and sex categories, habitat preference and distribution during both the wet and dry seasons. A line-transect census method was employed to assess the current population status of African elephants as adopted by Ratti *et al*. (1983); Megaze (2015) and Jyoti *et al*. (2020) for different mammals.

### 2.4. Population structure and estimate

Boundaries of zones were demarcated based on the understanding of the topographic and habitat types. Based on the number of transects, 41 patrolling teams were established, each composed of two-three Park/wildlife scouts, experts and trained Park staff. Daily monitoring data sheets, hybrid binoculars with range finder, compasses and radio for communication were also assigned to a team to collect the necessary information and each group assessed one transect a day. A total of 41 transects of different sizes were assessed to cover an estimated 25 percent of the total area in each habitat type. Training was given for the patrolling team on how to operate and use the GPS receiver, compass, binoculars, and data recordings in the field and how to record animal - observer distance, and perpendicular distances.

A total of 41 transects of different sizes were assessed to cover an estimated 25% of the total area in each habitat type. The length of transects ranged from 4.5 to 5 km, totaling a distance of 197 km. Eight repeated walks were made on transects 2 in each season during Year 2020-2021. Transect counts were carried out for each season both in 2020−2021 during wet and dry seasons from 06:00 to 10:00 h in the morning and 16:00 to 18:00 h in the late afternoon, when they were active and when visibility was good.. We walked a total of 788 km on the transects and recorded data on sightings of, number of individuals, group size, habitat types and age-sex structure of each individuals in a herd, animal-to-observer distance, angle of detection from main bearing and animal transect (perpendicular distances). We measured observer-to-animal distance using hybrid binoculars with rangefinders, and the angle of detection from the transect line using a compass. When elephants were encountered in a herd we recorded distance and angle to the centre of the herd.

Data were analyzed using DISTANCE version-7.4 software and computed estimate of density. We pooled the data from temporal replicates of each transect and treated the mean as a single sample and the mean cluster size was used for analysis. Variance in encounter rates of animals between transects was estimated empirically.

DISTANCE was selected the best possible model. We generated encounter rate, average probability of detection, cluster density, cluster size and animal density using the selected model in program DISTANCE. For sightings in which age-sex of individuals could not be recorded, we recorded only group size. Using a combination of characteristics such as differences in height, relative body size, external genitalia, size and shape of tusks and shape of the head profile (Demeke, 2010; Moss, 1996; Manspeizer and Delellegn, 1992). When first observing an elephant or groups of elephants in any locality, the sex and age structure and composition were recorded. Age classes were assigned based on the physical characteristics of the individual. Observations were made by ground surveys during 2020-2021. Data obtained were grouped into the following sex categories: Infants and calves of both sexes and juveniles of both sexes, sub-adult male, sub adult female, adult male and adult female (Moss, 1996). Elephants were grouped in to five age groups (Moss, 1996; Lee and Moss, 1995); Infants and Calf (< 2 year old), Juvenile (2 < X < 10 years old), Sub-adult (10 < X < 15) and Adults (> 15 years) (Williams, 2002; Demeke, 2010). Elephants grow throughout their life time (Hanks, 1979; Moss, 1996). The larger an elephant is therefore, the older its age. Body size comparison was performed relative to the height of adult female elephant in the group. Infants and calves in their first year can fit under their mother’s belly and when they drink water they submerge their faces. Juveniles pass under the throat and sub-adults possess height above juveniles but below the adult females.

Transects were surveyed systematically with the help of trained and experienced scouts during wet and dry seasons at a constant speed to maximize the probability of seeing all individuals on both sides of the transect (Norton-Griffiths, 1978; Jyoti *et al*., 2020). Whenever African elephants were located, the distance and sighting angle from transects, animal transect distance (perpendicular distance) observable activities and habitat types were recorded. Silent detection method was followed to minimize disturbances (Wilson *et al*., 1996). Repeated counting of the same herd was avoided by conducting the survey in all of the habitats within a single day and by using recognizable features or unusual features such as herd size, group composition and distinct individuals with deformities on tusk, tail and ear (Wilson *et al*., 1996). Thus, all herds were individually recognized.

The mean number of individuals observed per transect was pooled together using distance software, and extrapolated to estimate the population for the whole study area. Population density was estimated using this population estimate divided by the extent of the study area (Wilson *et al*., 1996 and Jyoti et al., 2020).

Herd size and composition were recorded by direct observations. During the field assessments, the composition of the elephant group was recorded as follows: Female group - one or more adult or sub-adult females and calves, with no males greater than 20 years old; bull group - two or more males, with ages greater than 20 years old in the absence of females; mixed group - a female group with one or more adult males; and lone bull - a single male elephant, with no other elephants around (Douglas-Hamilton, 1972; Moss, 1988 ; Demke, 2010).

### 2.7. Distribution and habitat association

The method of Norton-Griffiths (1978) was used to describe the wet and dry season distribution. By taking each group or individual sighting as scores with respect to habitat types and comparing their frequencies to the relative availability of vegetation type, it was possible to detect the utilization of vegetation type and the distribution of the African elephant.

## Results

A total of 74 sightings of elephants on average were recorded on line transects. The density of individuals was estimated to be 0.53 elephants per km^2^ (95% confidence interval) and the percent coefficient of variation of density of individuals was 46.83. The total population estimate was 756 individuals (Table 1).

**Table 1.** Density estimate for elephants in the Chebera Churechura National Park for the years 2020 and 2021.

| Parameter | Point Estimate | Standard Error | Percent Coef. of Variation | 95% Percent Confidence Interval |  |
| --- | --- | --- | --- | --- | --- |
| D | 0.53600 | 0.25100 | 46.83 | 0.21864 | 1.3140 |
| N | 756.00 | 354.02 | 46.83 | 308.00 | 1853.0 |

A total of 69 sightings of elephants on average was recorded on line transects during the wet season of 2020-2021. The density of individuals was estimated to be 0.48 elephants per km^2^ (95% confidence interval) and the percent coefficient of variation of density of individuals was 47.03. The total population estimate was 686 individuals (Table 2).

**Table 2.** Wet season density estimate for elephants in the Chebera Churechura National Park for the years 2020 and 2021.

| Parameter | Point Estimate | Standard Error | Percent Coef. of Variation | 95% Percent Confidence Interval |  |
| --- | --- | --- | --- | --- | --- |
| D | 0.48687 | 0.23029 | 47.30 | 0.19709 | 1.2027 |
| N | 686.00 | 324.47 | 47.30 | 278.00 | 1696.0 |

A total of 79 sightings of elephants on average was recorded on line transects. The density of individuals was estimated to be 0.58 elephants per km^2^ (95% confidence interval) and the percent coefficient of variation of density of individuals was 46.36. The total population estimate was 826 individuals (Table 3).

**Table 3.** Dry season density estimate for elephants in the Chebera Churechura National Park for the years 2020 and 2021.

| Parameter | Point Estimate | Standard Error | Percent Coef. of Variation | 95% Percent Confidence Interval |  |
| --- | --- | --- | --- | --- | --- |
| D | 0.58513 | 0.27171 | 46.36 | 0.19180 | 1.1749 |
| N | 826 | 383.57 | 46.36 | 338.00 | 2010.0 |

Age structure and sex ratio of the observed elephant population during wet and dry seasons for years 2020 and 2021 are given in Tables 4.

**Table 4.**
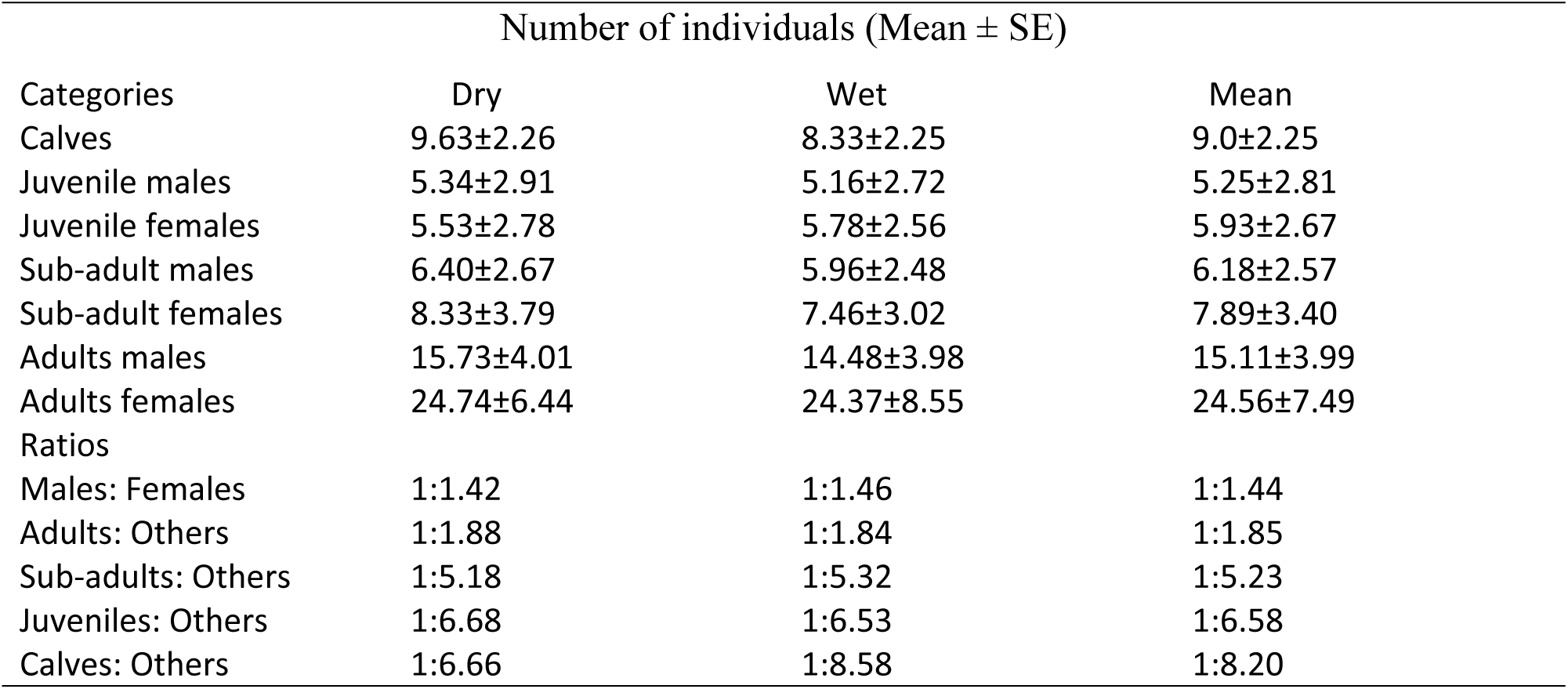
The mean age structure and sex ratio of the observed African elephant population during the wet and dry seasons for the years 2021 and 2022. Number of individuals (Mean ± SE)

| Number of individuals (Mean ± SE) | | | |
| --- | --- | --- | --- |
| Categories | Dry | Wet | Mean |
| Calves | 9.63±2.26 | 8.33±2.25 | 9.0±2.25 |
| Juvenile males | 5.34±2.91 | 5.16±2.72 | 5.25±2.81 |
| Juvenile females | 5.53±2.78 | 5.78±2.56 | 5.93±2.67 |
| Sub-adult males | 6.40±2.67 | 5.96±2.48 | 6.18±2.57 |
| Sub-adult females | 8.33±3.79 | 7.46±3.02 | 7.89±3.40 |
| Adults males | 15.73±4.01 | 14.48±3.98 | 15.11±3.99 |
| Adults females | 24.74±6.44 | 24.37±8.55 | 24.56±7.49 |
| Ratios |  |  |  |
| Males: Females | 1:1.42 | 1:1.46 | 1:1.44 |
| Adults: Others | 1:1.88 | 1:1.84 | 1:1.85 |
| Sub-adults: Others | 1:5.18 | 1:5.32 | 1:5.23 |
| Juveniles: Others | 1:6.68 | 1:6.53 | 1:6.58 |
| Calves: Others | 1:6.66 | 1:8.58 | 1:8.20 |

An average of 74 individuals of elephants was recorded during wet and dry season for the years 2020 and 2021. Among them, females constituted 52.12% and males 36.07% of the population (Table 4).

Mean of sex and age structure of elephants during wet and dry seasons for two years are given in Table 5. The mean of sex and age structure of elephants during wet and dry seasons for the years 2020 and 2021 are given in Table 5. The mean number of herd and the mean herd size of elephants during the wet and dry seasons for the years 2020 and 2021 are shown in (Table 5). In CCNP the number of herds observed and the number of individuals in each herd was different during the study period. A mean of 19 and 10 herds were recorded during the wet and dry seasons for the years 2020 and 2021respectively (Table 5). There was a significant difference in the mean number of herds (χ2 = 42.5, df =1, P<0.05). The herd size ranged from 1 to 149 individuals and the mean herd size during the wet and dry seasons were 16.5 and 50.25, respectively. There was a marked difference in the mean herd size between the two seasons (χ2 = 39.1, df =1, P<0.05).

**Table 5.**
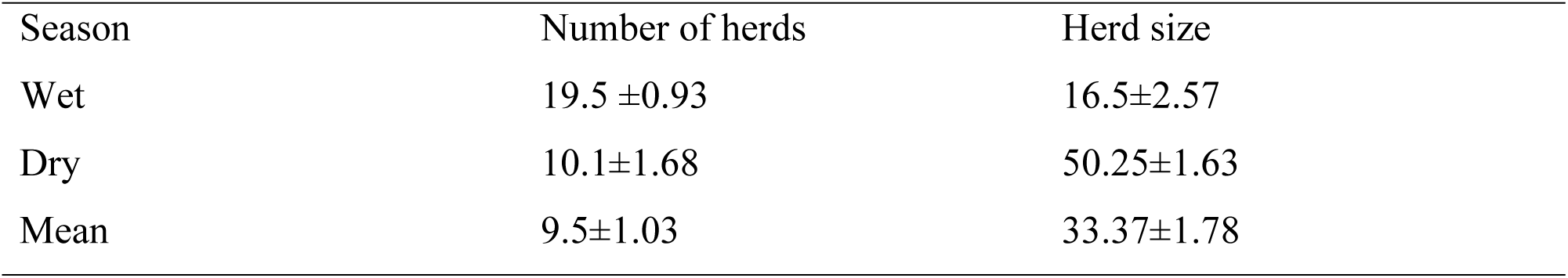
Number of herds and herd size of African elephants observed during the wet and dry seasons for the years 2020 and 2021 (Mean+SE) of the study period.

### 3.2. Distribution and habitat association

The relative use of different plants by the African elephant is indicated by the number of individuals observed in each vegetation communities (Table 6). Habitat selection varied seasonally. They showed high preference for riverine vegetation during the dry season. Out of the total 75.8% of the African elephants utilized riverine habitat during the dry season, whereas 21. 25% used this habitat during the wet season. Grassland with scattered tree habitat was utilized by 5% of the elephants during the wet season and only 3. 6%during the dry season. Woodland habitat was utilized by 71 % of the elephants during the wet season, and 17.65% during the dry season. Montane forest was utilized by 2.75% of the elephants during the wet season, and 2.95% during the dry season (Table 6). The distribution of African elephants in different habitat types during the wet and dry seasons showed highly significant variation (χ2 = 17.54, df =3, P<0.05).

**Table 6.** Observation of the African elephants in different habitat types in Chebera Churchura National Park during the wet and dry seasons. Number of individuals in different habitats Season

| Season | Number of individuals in different habitats Season |  |  |  |
| --- | --- | --- | --- | --- |
|  | GL | WL | MF | RF |
| Wet | 3.57 (5%) | 50.81 (71 %) | 1.94 (2.75%) | 15.2(21. 25%) |
| Dry | 2.72 (3. 6%) | 13.62 (17.65%) | 2.08(2.95%) | 57.76(75.8%) |
| Mean | 2.84 (4.53%) | 32.90 (44.32%) | 2.03(2.75%) | 36.04 (48.40%) |

GL= Wooded grassland, WL= Woodland, MF= Montane forest, RF= Riverine

## 4. Discussion

### 4.1. Population estimate of African elephants

Accurate and reliable recent information about the status, density and age-sex structure of African elephants for CCNP is lacking. Thus the present study can be a starting point and baseline for future studies in population trends and designing and implementing appropriate conservation programme and activities.

The current population estimate of African elephant (756 elephants, 0.53 elephants/ km^2^) is incomparable with the previous survey carried out by Admasu (2006) who estimated the total population at (84 individuals, 0.07 elephant km^2^). This might be due to differences in sampling methods utilized. Such a big difference in population estimate of the present study might be due to the extended period of time since the survey conducted and there is a considerable increase in the African elephant population in CCNP following the crackdown of ivory poaching after the establishment of the National Park in 2005 (Tsegaye, 2022). Moreover, there are also accuracy issues associated with the dung count method utilized, due to different factors such as; the time of counting which is usually during the summer/dry period due to logistics and convenience, and factors associated with two of the three parameters (dung defecation and dung decay rates) in the dung counting method. Researchers confirmed that there is up to 72% variation in dung decay rates in the same area between localities, seasons and years which causes biases in elephant density estimation (Kumara *et al*., 2012).

The African elephant population estimate in CCNP showed differences during wet and dry seasons. The dry season census showed high elephant population estimate compared to the wet season estimates. This was primarily due to the better quality fodder available during the dry season in the study area. The availability of lush growing grasses and green vegetation especially in the habitats adapted to fire such as wooded grassland and wood lands after the onset of wildfire during the dry season. Fresh grass and newly grown green leaves provide additional fodder in the wooded grassland and wood-land habitats (DeBoer *et al*., 2000; Admasu, 2006; Megaze, 2015 and Mekonnen, 2019). This was also confirmed with the study of Sinclair (1977) in Serengeti National Park of Tanzania. Burning practice of the area is found to be one of the main factors influencing the movements and distribution of African elephants and other wild animals especially during the dry season. Even though breeding in most elephant populations does not exhibit a pronounced seasonality the occurrence of oestrus and conception is sensitive to rainfall and resource availability (Laws and Parker, 1968; Laws, 1969; Poole, 1987 and Moss, 1988).

According to Poole (1987) and Moss (1988), estrous females may be observed in any month of the year. The frequency of oestrus is significantly higher during and following the wet seasons in the Amboseli population when females are in good condition. The seasonality of male musth periods also reflects the pattern exhibited by females. The result of the present study showed that unlike the points mentioned above due to the unique and prolonged rainy season in CCNP which lasts about nine months and the early setting of management and wild fire in most of the woodland and grassland habitats from early November to January new growth of fresh grass and green leaves triggered by the fire provides additional better quality fodder in those habitats types in addition to the riparian forest during late dry season from March to May (DeBoer *et al*., 2000; Megaze, 2015; Tsegaye *et al*., 2015). Moreover, the annual fire in most of the woodland and grassland habitats makes the habitats open and there was better condition for counting animals during the dry season (DeBoer et al., 2000; Megaze, 2015; Tsegaye *et al*., 2015; Mekonenne, 2019). As a result more young animals (calves and yearlings) were observed during this season which may be explained by the findings of the present study which showed more increased population of elephants during the dry season than the wet season. High proportion of female to male ratio was recorded in the population indicating that elephants have a potential to increase in number. This may be due to male and old elephants were selectively poached for their large sized tusks before the crackdown of ivory poaching following the establishment of the Park. Such killings have a series consequence on the mating success of bulls since the oldest males; have better chance of succeeding in mating than younger ones. The result of the present study goes in line with the findings of Admasu (2006) and Kumara1 *et al*. (2012) who mentioned that the sex ratio of elephant population was dominated by female indicating that selective poaching on old and male elephants had been carried out for their large tasks in CCNP, Ethiopia and in Biligiri Rangaswamy Temple Tiger Reserve, India respectively.

Low proportion of young to that of other age groups was observed during the present investigation. This might be due to high mortality rate of young and negatively biased counting due to their small size and detectability. The finding of the present study goes in line with the findings of Admasu (2006) and Megaze (2013) who noted that that the presence of low proportion of young to that of other age groups was due to young ones are more vulnerable to predators and they are usually hidden under the dense grasses and riverine vegetation during the wet and dry seasons which might negatively affect their counting and detectability (Megaze, 2015; Tsegaye *et al*., 2015).

In CCNP, elephants were observed in smaller herds during the wet season and in larger herds during the dry season. The seasonal change of group size might be due to both changes in the availability of resources and their social behaviour as noted by Moss (1988) and Poole and Moss (1988) who mentioned that where resources are both plentiful and evenly distributed, elephants tend to aggregate and this is particularly noticeable in many places during and following the rains when resources are plentiful (Douglas-Hamilton, 1972; Leuthold, 1976; Western and Lindsay, 1984; Moss, 1988; Poole and Moss, 1989).

According to Moss (1977), as the dry season progresses the large aggregations and bond groups of elephants begin to split. A number of explanation has been proposed for these large aggregations including access to mates (Moss, 1988; Poole and Moss, 1989) and the renewing of social bonds (Moss, 1988). The finding of the present study goes in line with the above result showing large aggregations and bond groups of elephants during the late dry season when resources were plentiful due to the unique and prolonged rainy season in CCNP. Plenty of fresh food was found to be available during this season from March to May after the early setting of fire in most of the woodland and grassland habitats from November to January (Megaze, 2015; Tsegaye *et al*., 2015).

During this season following the availability of abundant resources in CCNP, large aggregations of elephants were recorded which may also be additionally influenced by social factors such as access to mates and the renewing of social bonds (Moss, 1988).

According to Laws (1969) and Poole (1987), the occurrence of oestrus and conception is related to rainfall and resource availability. Similar findings were also recorded in the Amboseli population. The occurrence of oestrous females was significantly higher during and following the wet seasons when females were in good condition (Poole, 1987; Moss, 1988).

The seasonality of male musth periods also reflected the pattern exhibited by females (Poole, 1987). Moreover, availability of permanent water sources and fruits of different tree species in the riparian forest habitat such as *Ficus* and *Phonex* within their dry season home range were also among the driving factors for such aggregations. In contrast to this in CCNP during the main rainy season fruits and other dry season food sources were not available to the elephant herds. Elephants will be forced to change their food sources to some herbaceous and few woody plant species in the wooded grassland and woodland habitats during the main rainy season. The annual grass that is newly grown during the dry season from early November to January has become very tall, dry and less edible due to the age effect (Megaze, 2015; Tsegaye *et al*., 2015).

The annual grass grown early in November and on which elephants depended upon will be very tall, dry, less edible and low in nutritive value during the main rainy season due to the growth factor but many annual water sources were available everywhere. Therefore, during this time, elephants disperse and form smaller groups for intensive foraging of the available resources. This was similar to the finding of Megaze (2015) and Tsegaye *et al*. (2015) for different animals. During the dry season, forest fires were common in CCNP, and there were new growth of a better quality grass due to the unique rainfall pattern in the area. Open and free spaces were also available during this season and large herd size of waterbucks and buffalos were recorded. In contrast to this, waterbucks and buffalos were dispersed and formed smaller herds for intensive foraging of the available edible food resources during the wet season (Megaze, 2015; Tsegaye *et al*., 2015).

### 5.2. Distribution and Habitat Association

Comparison of the seasonal changes in habitat association shows that elephants have high preference for riverine vegetation during the dry season and grassland vegetation during the wet season. During the wet season, African elephants were observed more in the wooded grassland habitats. They gradually moved away from the main riverine vegetation as the area becomes wet. The flooded riverine habitat and availability of alternative food and water sources may force elephants to move out of the riparian forest to the woodland and wooded grassland habitats that had previously been burn during the dry season. It is also less flooded, with re-growth of some lush green sward, at the intermediate stage of growth (Megaze 2015; Tsegaye *et al*., 2015; Mekonenne, 2019) Therefore, selection for wooded grassland habitat during the wet season coincide with increased need to avoid the already utilized and flooded dry season riparian forest habitat and move in to the relatively productive woodland and wooded grass land habitats covered with older annual grasses but with plenty of green woody plant species fully recovered from the annual fire. But, during the dry season starting from November to May clumped distribution of elephants occurs in the riverine vegetation. The association of elephants with the riverine vegetation increases significantly from the end of the rainy season as the dry season progresses. In addition, very tall, dry, less edible and low in nutritive value grass will persist in the wood land and wooded grassland habitats until the onset of fire during November and instead new growth of grass and recovery of green wood land tree species occur (Megaze. 2015; Tsegaye *et al*., 2015). The same finding was also confirmed with the study of Sinclair (1977) in Serengeti National Park of Tanzania. Burning practice of the area is found to be one of the main factors influencing the movements and distribution of African elephants and other wild animals especially during the dry season.

Therefore, only few numbers of elephants were observed in woodland and wooded grassland habitats during the dry season. The optimum habitat (the highest positive association) for elephants during the dry season was the riverine vegetation in the present study area. During the wet season previously avoided vegetation type were utilized to a greater extent as they become more suitable feeding habitat as a result of food scarcity due to rain and flooding in the already utilized riparian forest habitats (Megaze, 2015; Tsegaye *et al*., 2015).

Scattered distribution was observed particularly when the rain was evenly distributed in the Park. The present study shows that African elephants showed random association with the four habitat types (montane, riverine, woodland and wooded grassland forest habitats) throughout the study periods. Their distribution was not uniform across the four main habitats in the study area and showed differences due to the availability of food and permanent water sources.

## ACKNOWLEDGEMENTS

The authors greatly acknowledge Addis Ababa University for their logistic and research financing support. Moreover, the authors wish to thank staff members of Chebra Churchura National Park, to the local people of Dawro and Konta Zones, and their local administrators for their kind cooperation, patience, hospitality, and willingness to share their knowledge.

